# TiPS: rapidly simulating trajectories and phylogenies from compartmental models

**DOI:** 10.1101/2020.11.09.373795

**Authors:** Gonché Danesh, Emma Saulnier, Olivier Gascuel, Marc Choisy, Samuel Alizon

## Abstract

We introduce `TiPS`, an R package to generate trajectories and phylogenetic trees associated with a compartmental model. Trajectories are simulated using Gillespie’s exact or approximate stochastic simulation algorithm, or a newly-proposed mixed version of the two. Phylogenetic trees are simulated from a trajectory under a backwards-in-time approach (i.e. coalescent). `TiPS` is based on the `Rcpp` package, allowing to combine the flexibility of R for model definition and the speed of C++ for simulations execution. The model is defined in R with a set of reactions, which allow capturing heterogeneity in life cycles or any sort of population structure. `TiPS` converts the model into C++ code and compiles it into a simulator that is interfaced in R *via* a function. Furthermore, the package allows defining time periods in which the model’s parameters can take different values. This package is particularly well suited for population genetics and phylodynamics studies that need to generate a large number of phylogenies used for population dynamics studies. This package is available on the CRAN at https://cran.r-project.org/package=TiPS.

## Introduction

Stochastic population dynamics simulations are routinely used in ecology and epidemiology to generate trajectories (i.e. time series of population sizes) and genealogies that capture the relatedness between individuals [1–3]. The increasing amount of genetic data is fuelling interest in linking population dynamics and genealogies because the former can leave footprints in individuals’ genomes [4–6]. Such phylodynamics studies involve computer-intensive methods that can require the simulation of many trajectories and genealogies [7–9].

A common method to simulate population dynamics trajectories is Gillespie’s exact stochastic simulation algorithm (SSA) [10], which is rooted in probability theory [11]. In the R software environment, it is implemented in packages such as `Gillespi`eSSA [12], `adaptivetau` [13], or `epimdr` [14]. In this same environment, `geiger` [15], `phytools` [16], `ape` [17], and `TreeSim` can simulate phylogenies using a birth-death model. However, few software packages simulate both trajectories and genealogies. One exception is `rcolgem`, updated to `phydynR`, in R [18], which involves coalescent processes. Another exception is the software package `MASTER` [19] in the BEAST2 platform [20].

We introduce `TiPS`, a flexible and easy-to-use R package to rapidly simulate population trajectories and phylogenies using a backwards-in-time, i.e. coalescent, process with either pre-defined sampling dates or a stochastic sampling scheme. We also introduce a new approximate version of the Gillespie algorithm to increase calculation speed. A brief benchmarking analysis shows that `TiPS` is faster than `adaptivetau` to simulate trajectories, especially for large populations (Figure 3a). It is also at least one order of magnitude faster than `phydynR` to simulate phylogenies (Figure 3b).

## Methods

### Package overview

`TiPS` generates two types of stochastic simulation outputs: population dynamics trajectories and phylogenies. These are obtained using a continuous-time individual-based model defined in R as a system of reactions. The model is first transcribed in C++ and then compiled, before being linked back to a simulating function in R thanks to the `Rcpp` package [21]. The general structure of the pipeline is illustrated in Figure 1.

**Figure 1:**
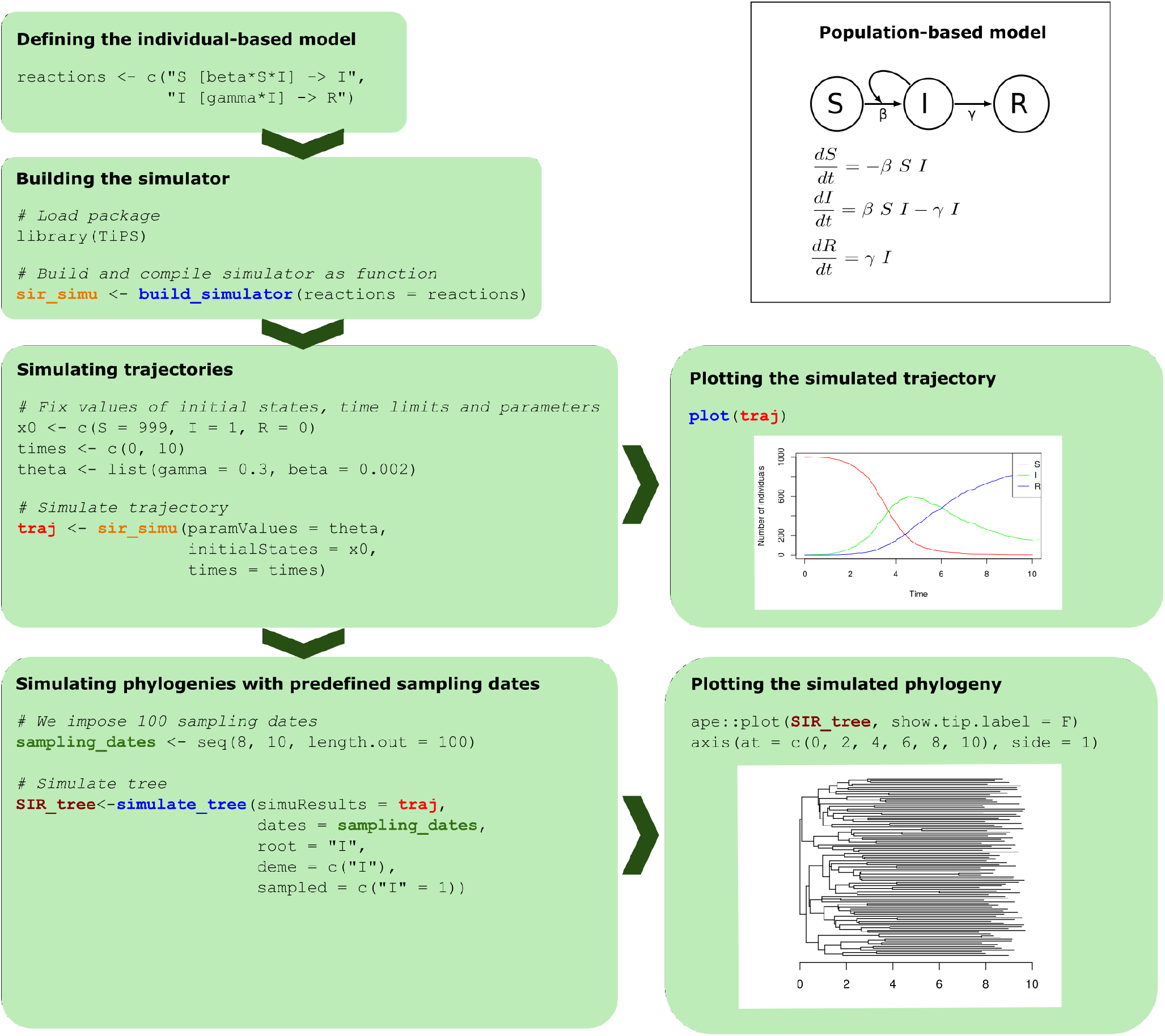
General structure of the TiPS pipeline. The equations and outputs correspond to the *SIR* epidemiological model [2]. The functions of the R package are in blue. The simulator of trajectories, which is built as a function, is in orange. The variable *traj*, in red, is the output trajectory of class *simutraj*. It can be plotted using our *plot* method. `TiPS` used the simulated trajectory and 100 sampling dates that we generated (variable *sampling_dates* in green) to simulate a sampled phylogeny. The output simulated phylogeny is a *phylo* class object from the `ape` R package [17], which can be used for plotting.

### Model description

`TiPS` simulates trajectories from compartmental models. These models divide the population (animals, cells, *etc*) into distinct categories (geographic, clinical state, *etc*) or so-called compartments in which the sub-population behaves uniformly. The population, in these models, may progress between the different compartments.

Here, we illustrate the use of `TiPS` with the SIR epidemiological model, where individuals can have three clinical states: susceptible (*S*), infected (*I*), and removed (*R*) [2]. The model can be captured with a system of two individual-based reactions:

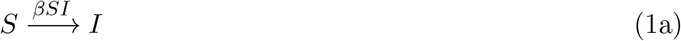

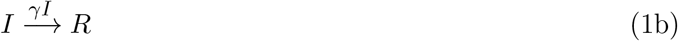

where *β* and *γ* are the transmission and recovery rates. The rate of occurrence of each reaction is indicated above the transition arrow and the corresponding population-based system of ordinary differential equations (ODE) is shown in the top-right panel of Figure 1.

A simulating function is generated from the individual-based reactions of the model of interest. These are entered by the user as a string vector (top left box of Figure 1).

### Simulating trajectories

The simulation function takes as arguments a named numeric vector that contains the initial number of individuals in each compartment, a named list with the parameter values, a vector of the time limits of the simulation, and the type of algorithm to use for the simulation. Users can also enter a vector of breakpoints, which allows parameter values to vary over time. These breakpoints are indicated in the time limits vector, and the corresponding parameter values are ordered chronologically in the named list of parameter values.

Three simulation algorithms are implemented in the `Rcpp` simulating function:

1. Gillespie’s Direct Algorithm (GDA, the default option) [10] is an exact algorithm that simulates the time until the next event *δ*_*t*_ by assuming that waiting times are exponentially distributed. A limitation is that its computational complexity scales linearly with the number of simulated events.
2. Gillespie’s Tau-Leap Algorithm (GTA) [22] is an approximate algorithm that introduces a fixed time step *τ* during which the number of events of each type is assumed to be Poisson distributed. This algorithm is limited in terms of computation time if the time step is small compared to the rate at which events occur.
3. The Mixed Simulation Algorithm (MSA) is a new algorithm we introduce that switches from GDA to GTA depending on the respective values of *δ*_*t*_ and *τ*. The algorithm switches from GDA to GTA if 10 successive estimations of *δ*_*t*_ are shorter than *τ/*10. The algorithm switches back to GDA if the total number of realised events is lower than the number of possible events. The MSA shares similarities with the slow-scale stochastic simulation algorithm [23] or the adaptive explicit-implicit tau-leaping method for an optimised tau-leap selection [24].

The output of a trajectory simulation is a named list containing the simulated trajectory and some details about the model and the simulation, such as the reactions of the model, the parameter values, the time range, the algorithm used to perform simulations, the time-step in case the algorithm is the GTA or the MSA, and, for reproducibility purposes, the random seed used to initialise a pseudorandom number generator.

### Simulating phylogenies

A phylogeny is the representation of the evolutionary history and relationships between genes, organisms, or groups of organisms. The root of the tree represents the ancestor of all lineages, and the leaves represent the most recent descendants of that ancestor. `TiPS` simulates binary phylogenies that are rooted in time. We distinguish the biological process from the observation process. The former corresponds to a sequence of individual birth, migration, and death events. The latter corresponds to sampling or other events that may interrupt the biological process (e.g. sterilisation or treatment).

`TiPS` uses a coalescent approach [25] to simulate phylogenetic trees based on trajectories, which correspond to a list of dated events (or ‘reactions’) and sampling dates (e.g. based on observed data). Each node in the simulated phylogenetic tree is associated with a state, i.e. a compartment name, and its height, i.e. its distance to the root.

Individuals who can contribute directly to the phylogeny are referred to as ‘demes’ and the compartment to which they belong as a ‘deme compartment’. Individuals in non-deme compartments are not observed in the phylogenetic process. For infection phylodynamics, when simulating a phylogenetic tree, we trace back the history of sampled infections. In the SIR model, for example, these are only found in the I compartment (S and R are non-deme compartments). In an ecological context, under a predatorprey model, for example, tracking the history of predator and prey individuals means that both are deme compartments.

`TiPS` can simulate the phylogeny of the full trajectory or that of a sampled phylogeny if sampling dates are provided. These dates can either be provided by the user as a vector (bottom left panel of Figure 1) or generated during the trajectory simulation by adding a sampling reaction in the model. Since parameter values can vary with time, `TiPS` can reproduce temporal variations in sampling intensity. If sampled individuals can belong to more than one deme compartment, the user can choose between defining the state associated with each sampling event or defining a proportion of sampling events associated with each state. In the last case, `TiPS` randomly associates a state to each sampling date.

After this pre-processing, the sampling dates are organised as a named list containing a vector of decimal dates (column name ‘Date’) and a vector containing the reactions indicating the state of individuals to sample (column name ‘Reaction’). `TiPS` then incorporates these pieces of information into the recorded trajectory (containing also dates and reactions) in chronological order.

The simulation of a phylogeny (sampled or not) starts from the most recent sampling date (or the most recent death event) and progresses through the simulated trajectory backwards-in-time. When simulating a phylogenetic tree, each event in the trajectory is defined as one of the following types of tree reactions:

- births: these reactions correspond to the creation of a new deme individual;
- migrations: these correspond to the displacement of a deme individual from its deme compartment to another;
- death: these correspond to the removal of a deme individual from its deme compartment or the displacement of a deme individual from its deme compartment to a non-deme compartment;
- samplings: these correspond to the sampling of deme individuals.

Table 1 illustrates demes and reaction types for specific ecological and epidemiological models.

**Table 1:**
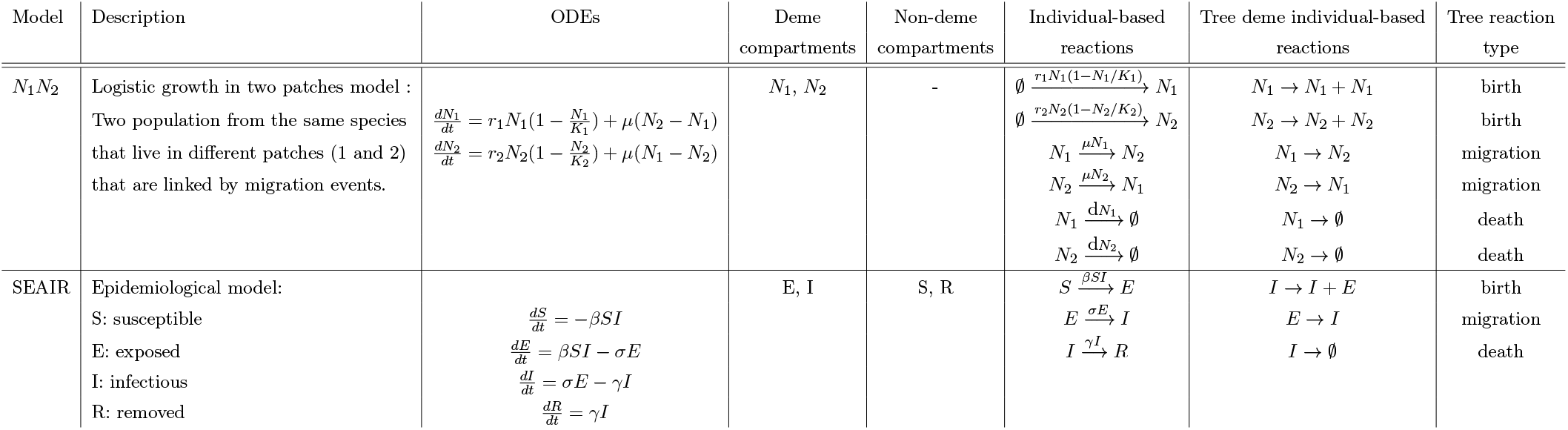
Description of demes and reactions for an ecological and an epidemiological model. ‘ODEs’ stands for ordinary differential equations.

Each of these four types of reactions can result in a modification in the simulated tree. A forward-in-time birth event can be represented by a coalescence between two lineages under a backwards-in-time process (or a branching under a forward-in-time process). A sampling event interrupting a biological process is represented as an external node (or leaf) in the phylogeny, and a death event, if observed, is also represented as a leaf. A migration event of an individual from one deme compartment to another will change and update the state of its corresponding lineage.

In some cases, especially with the tau-leap method, more than one event may occur on the same date (e.g. three new births). To determine the number of events that lead to a change in the phylogeny (e.g. a branching), we draw a number from a hypergeometric distribution, which is appropriate since it describes the number of successful events (*k*) when drawing *n* times (without replacement) from a total population of size *N* that contains *K* elements corresponding to a ‘success’. For example, when a birth event in the trajectory is encountered in the phylogeny simulation, the algorithm draws, using the hypergeometric distribution, the number of sampled child lineages and the number of sampled parent lineages to coalesce into one sampled parent lineage. If at time *t*, the number of sampled individuals is small, all birth events at time *t* may not lead to an observed branching in the tree. Similarly, when a migration event is encountered, the algorithm draws the number of sampled lineages associated with a change in state. Figure 2 illustrates how the tree is simulated from the trajectory and further details can be found in the Appendix.

**Figure 2:**
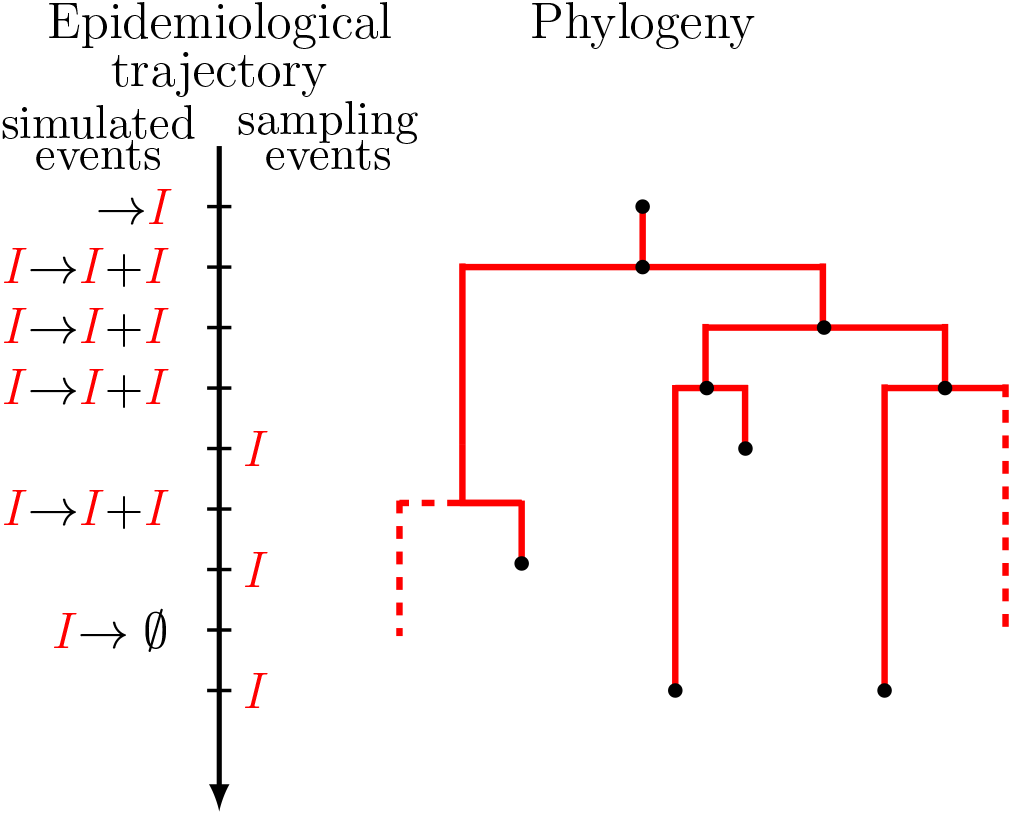
Tree simulation for a SIR model. the trajectory shows the series of epidemiological events and sampling events. the phylogeny represents the epidemiological history of the individuals carrying sampled viruses in solid lines. The rest of the history is shown in dashed lines. In this representation, at each birth event (branching), the donor is deviated to the right side and the recipient to the left side.

The output simulated phylogeny is an R object of class *phylo* as defined in the `ape` package [17].

An illustration of how to simulate a phylogeny can be found in the ‘Simulating phylogenies’ box in Figure 1 and in the Appendix for a logistic growth ecological model.

### Benchmarking

To evaluate `TiPS`’s performances, we designed a benchmarking analysis on both modules of the software package, i.e. the trajectory and the phylogeny simulators, comparing with existing `R` packages (`adaptivetau` and `phydynR`). Table S1 summarises the main features of the approaches used.

We first evaluated the computational speed and the accuracy of the trajectories (i.e. populating dynamics). For five initial population sizes and three different `R` packages, we simulated 10, 000 trajectories of the epidemiological Susceptible-Infected-Recovered (SIR) model and measured the execution time.

We then compared the time to simulate phylogenies under an SIR model. We varied sampling proportions in order to obtain target phylogenies of various sizes (10, 100, 500, 1000, and 1500 leaves). We simulated 1,000 phylogenies under each sampling scheme using `phydynR` and our package.

Furthermore, assuming a more detailed epidemiological model with two host types, as described in [26], we simulated an epidemiological trajectory using the tau-leap algorithm, and a complete phylogeny, such that each end of infection event corresponded to a leaf in the tree. The simulation generated a full transmission chain corresponding to a phylogeny with 154, 507 leaves. From this complete phylogeny, we generated 10 sub-trees by randomly sampling and keeping 1, 000 leaves for each sub-tree. We then used `TiPS` and `phydynR` to simulate 1, 000 phylogenies with each package under the same epidemiological model with the same parameter values. Note that the 1, 000 dates of each target sub-tree were imposed when simulating these phylogenies using a backwards-in-time approach. To compare the 2, 000 simulated phylogenies with the target one, we computed 60 summary statistics for each of them using the methods described in [9].

## Results

In Figure 3(a), we show the median execution time for one trajectory simulation and the 50% interquartile envelope. `TiPS` and `adaptivetau` rely on Gillespie’s Direct method (GDA), whereas `phydynR` uses the Euler-Maruyama integration (EMI). As expected, the population size, and, hence, the number of events per unit of time, increases the execution time for GDA-based packages but not for the EMI-based package. However, `TiPS` remains faster than the other two software packages for large populations (10^7^ individuals). In Figure 3(b), we perform the same simulations using approximations of the Gillespie algorithm with fixed time steps. The computational speed of this GTA implemented in `TiPS` and `adaptivetau` is comparable and much faster than the GDA. Our new MSA algorithm outperforms existing methods and improves computational speed compared to our GTA, especially for small population sizes.

**Figure 3:**
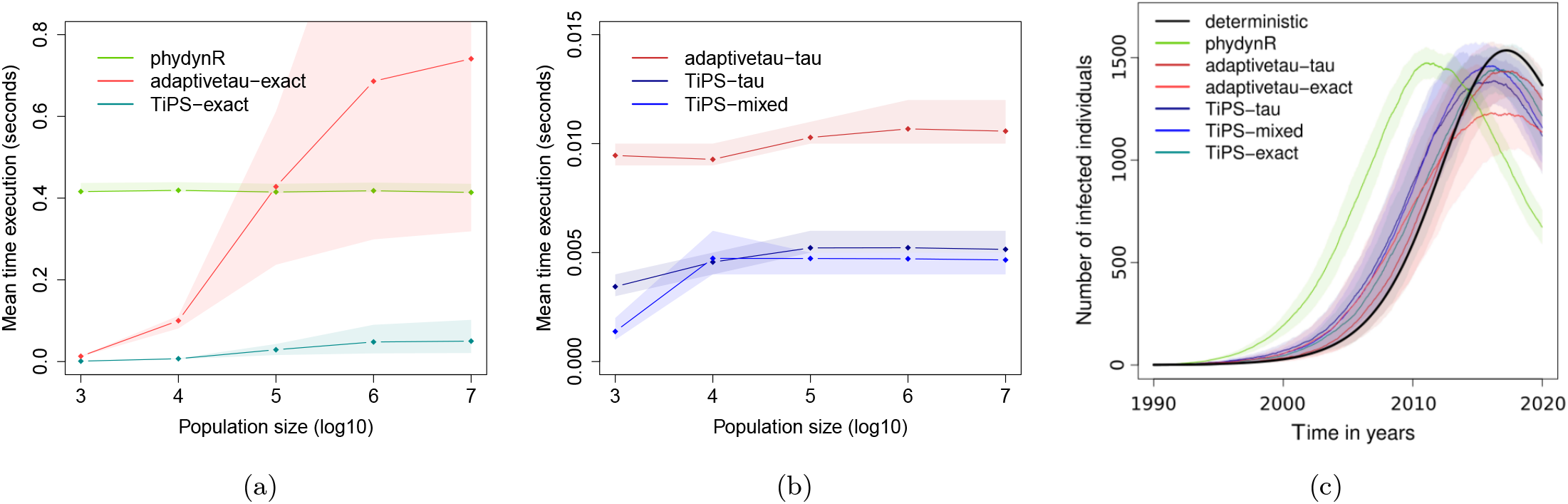
Benchmark analyses of trajectories simulations. a) Median computation speed and 50% confidence intervals (CI) for one simulation using GDA or EMI methods, b) median computation speed and 50% CI for one simulation using approximating GDA methods, and c) mean resulting trajectories for prevalence time series and the 90% CI. GDA stands for Gillespie’s Direct Method and EMI for Euler-Maruyama Integration. 10, 000 trajectories simulations were performed under an epidemiological SIR model, for five initial population sizes *N* varying from 10^3^ to 10^7^ and with parameter values *ℛ* _0_ = 2, *γ* = 1/3 and 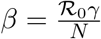.

In Figure 3(c), we show the deterministic trajectory, and, for each algorithm used, the mean simulated trajectory and its 90% confidence envelope. Among the packages studied, `TiPS` (in blue) is the one that yields the trajectories that are the closest to the deterministic prediction (in black). Note that there is a temporal shift for all the stochastic simulations, with a more rapid increase in population size compared to the deterministic model. This comes from the fact that stochasticity tends to disfavour trajectories that spread more slowly than average because they are more likely to go extinct. This known population genetics effect has been described in epidemiology models [27].

We then analyse the median time executions of simulations of phylogenies for each tool and each sampling scheme (Figure 4(a)). `TiPS`’s median simulation time is several orders of magnitude faster than that of the other method, with a more pronounced advantage for large phylogenies.

**Figure 4:**
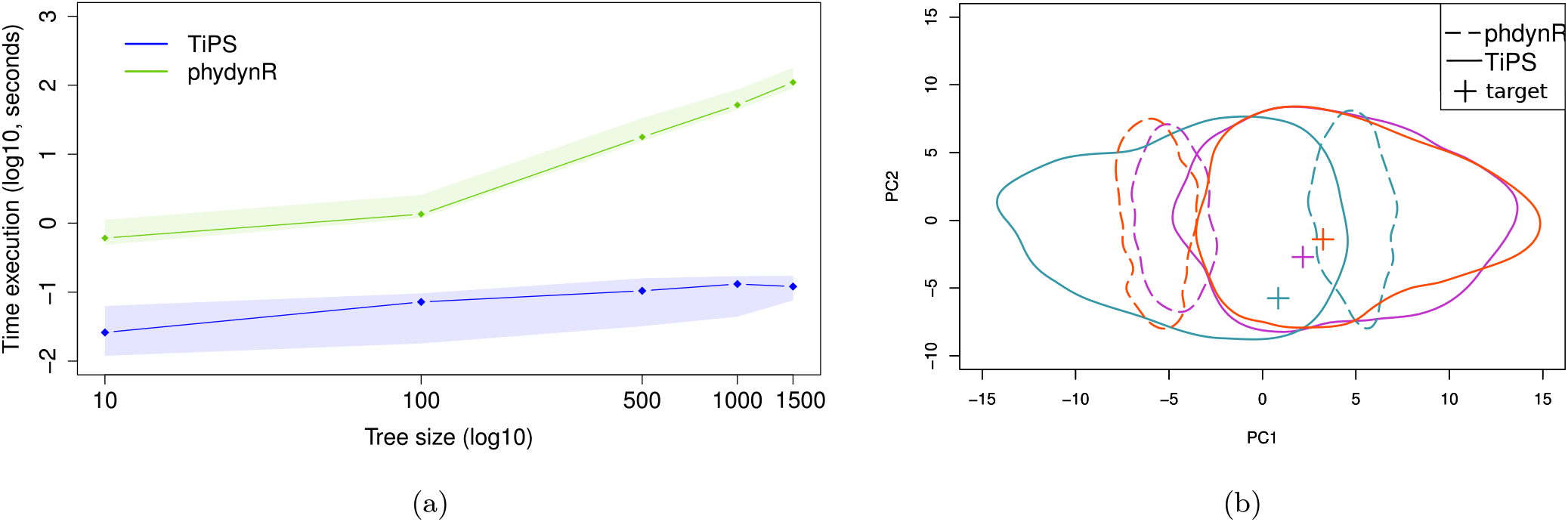
Benchmarking analysis of phylogenies simulations. a) Median computation speed for simulating one phylogeny of a given size and b) First two axes of a Principal Component Analysis (PCA) of phylogenies summary statistics. In a, the shaded area shows the 95% confidence intervals. In b, the cross shows the target phylogeny. The three colors represent three analyses using different target sub-trees. The phylogenies summary statistics used for the PCA are the same as in [9].

Regarding the accuracy of the simulations, we see in Figure 4(b) that the cross, which indicates the projection of the summary statistics from the target phylogeny, is contained in the envelope containing 90% of the phylogenies simulated using `TiPS` but not of those simulated using `phydynR`, and that for three different target phylogenies used represented by distinct colors. We observed the same results for seven other target phylogenies used. This means that the target phylogeny cannot be distinguished from phylogenies simulated using the same model with the same parameters using our package. The discrepancy observed for `phydynR` could originate from the trajectory, which strongly differs from the deterministic prediction (Figure 3c). Furthermore, `TiPS` is favoured in this analysis because the same algorithm (Gillespie’s Tau-Leap) was used to simulate the target phylogeny and the 1, 000 sub-trees. Supplementary Figure S7 shows the distributions of summary statistics computed from the phylogenies simulated using `TiPS` and `phydynR` for each analysis using a different target tree.

## Discussion

We developed a flexible R package to rapidly simulate trajectories and phylogenies from compartmental models. Its structure allows the user to include several sources of heterogeneity between individuals or between populations, e.g. different life stages or metapopulation structures. The simulation of the phylogeny relies on the trajectory and involves a coalescent approach.

Our benchmarking analyses show that `TiPS` is comparable to or outperforms existing `R` packages in terms of speed when generating numerous trajectories or phylogenies. The same is true for the accuracy of the simulation outputs.

The first asset of this software package is its flexibility since it can readily be used for any compartmental model. A second asset is its computation speed as it can simulate trajectories in a matter of milliseconds on a regular desktop computer. These properties have already been used for infection phylodynamics studies involving Approximate Bayesian Computing (ABC) [26] or to illustrate the effect of superspreading events [28].

Beyond epidemiology, this software package can be used more broadly to simulate population dynamics and associated genealogies. Some future extensions of `TiPS` will consist in introducing non-Markovian dynamics, simulating multifurcating phylogenies and implement other optimised stochastic simulation algorithms [23, 24].

## Supplementary Information

### S1 Phylogenies simulation algorithm

`TiPS` simulates rooted and binary phylogenies, meaning that every internal node has exactly two daughter nodes. The root of the tree represents the ancestor of all lineages, and the tips of the branches (or leaves) represent the most recent descendants of that ancestor. A coalescence between two lineages using a backwards-in-time process is equivalent to a branching event in the phylogeny in the forward-in-time process. Under the hypothesis of neutral evolution that phylodynamics relies on, a branching event can represent, for example, a viral transmission event, and a leaf can represent the end of viral infection. Note that we will refer to a node’s height as its distance to the root.

In this section, we distinguish two types of compartments: the deme compartments, where individuals contribute directly or indirectly to the phylogeny, and the non-deme compartments that cannot be placed in the genealogical process. For example, in a SIR epidemiological model, the deme and non-deme compartments would be respectively the I compartment and the S and R compartments. Each deme compartment is denoted *X*_*i*_, with *i* ranging from 1 to the number of deme compartments in the model. Sampled individuals belong to the sub-compartment 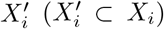 and are all associated with a leaf in the tree. We also introduce 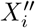, the sub-compartment of 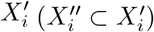 corresponding to individuals in 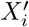 that have not yet been but may be sampled in a backwards-in-time process. The discrete size of compartments 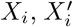 and 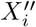 at time t are denoted as 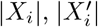 and 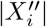, respectively. A non-deme compartment is denoted *Z* and its discrete size |*Z*|.

The tree simulation starts from the last (i.e. most recent) sampling date and progresses through the simulated trajectory backwards-in-time. The number of events that lead to a change in the phylogeny is drawn from a hypergeometric distribution. The hypergeometric distribution is appropriate as it describes the number of events *k* from a sample *n* drawn from a total population *N* without replacement. Each of the four types of reactions (sampling, birth, death, and migration) can result in a modification in the simulated tree : a new external node (or leaf) or the coalescence of two lineages.

#### Sampling event

We define a sampling event as an event that interrupts the biological process of an individual in the population of interest, and a re-sampling event as an observation event. In an ecological context, the sterilisation of an animal would be a sampling event whereas marking the animal would be a re-sampling event. In an epidemiological context, that is when tracing back the history of pathogen spread, a sampling event usually corresponds to a host individual being detected. Usually, this is assumed to coincide with the end of the infectious period, because the individual will be isolated or use adequate protection to prevent further transmission. However, if this assumption does not apply, it is possible to treat the event as a re-sampling event, such that the host individual will continue to transmit the pathogen to other individuals.

When we know the sampling dates, we know the number of sampling events occurring at time 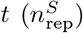 and, therefore, the number of tips to create with height *t*. However, since we allow sampled lineages to continue to have an offspring, we need to determine if the nodes we sampled at time *t* are associated with re-sampling events or not (i.e. if they should be linked to a node that has already been sampled after time *t* in our coalescent approach). The number of re-sampling (*n*_*RS*_) and sampling events (*n*_*S*_) at time *t* is governed by the following relationships:

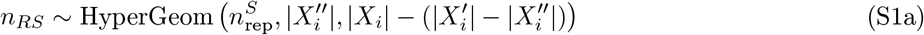

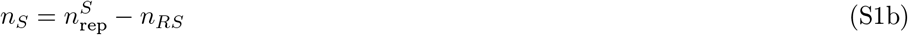

where 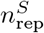 is the total number of phylogeny nodes generated at *t*. Using the hypergeometric distribution, we compute the number of re-sampling events *n*_res_ if we sample 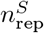 times without replacement in a sample of size 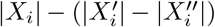 containing 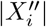 individuals. The number of ‘classical’ samplings, *n*_*S*_, is the difference between *n*_RS_ and the number of tips to be generated 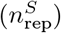.

In the case of a re-sampling, we randomly pick a node from 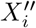 (the pool of individuals who have not yet been sampled), update its height to *t*, and link it to a node from 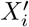 (the pool that has been sampled). Each re-sampling event decreases 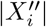 by one. In the case of a ‘classical’ sampling event, a new node with height *t* is created in 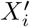 and 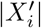 increases by one.

#### Birth event

A birth reaction can be written as follows: *X*_*i*_ → *X*_*j*_ + *X*_*i*_ where *i* and *j* range from 1 to the number of deme compartments. The birth reaction can also be written as *Z* + *X*_*i*_ → *X*_*j*_ + *X*_*i*_ if it includes a non-deme individual. In this section, we will refer to a donor lineage as the parent lineage and a recipient lineage as the child lineage. A birth reaction leads to one of two types of modification in the tree: a coalescence or an ‘invisible’ coalescence. A coalescence corresponds to the coalescence of two sampled lineages from two individuals, the recipient (or child) and the donor (or parent), into one sampled individual lineage (the donor). An ‘invisible’ coalescence is defined as the coalescence of the sampled lineage of the recipient 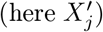 and an unsampled lineage of the donor 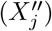 into one unsampled lineage 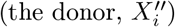.

We first need to determine the number of recipient lineages *n*_recipient_. Again, assuming a hypergeometric distribution, we compute the number of recipient lineages *n*_recipient_ if we sample *n*_rep_ times without replacement in a sample of size |*X*_*j*_| containing 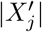 individuals (see equation S2a). If there are no recipient lineages (*n*_recipient_ = 0), there is no coalescence or invisible coalescence and, therefore, no change in the tree. Otherwise, we need to determine the number of donor lineages *n*_donor_. Since the sampling is performed without replacement, the total sample size is updated to |*X*_*j*_| − *n*_rep_ and the number of daughter lineages *Y* represented by nodes that are available for the coalescence becomes 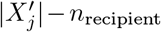. Thus, we compute the number of donor lineages *n*_donor_ if we sample *n*_rep_ times without replacement in a sample of size |*X*_*j*_| − *n*_rep_ containing 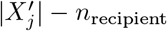 individuals with *i* = *j* S2b, or in a sample of size |*X*_*j*_| containing 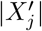 individuals with *i* ≠ *j* S2c. Mathematically, we can write:

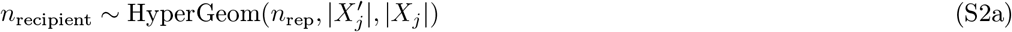

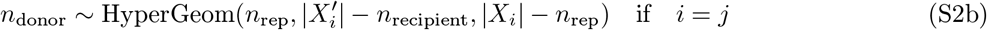

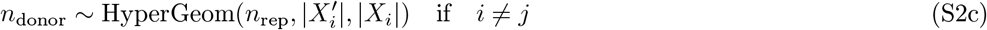

Knowing the number of recipient lineages, we need to determine the number of coalescences (*n*_*C*_), *i*.*e*. the number of lineages among the *n*_donor_ sampled donor lineages that will coalesce with the recipient ones. The number of visible coalescences is drawn from a hypergeometric law where we sample *n*_recipient_ times in a sample of total size *n*_rep_ containing *n*_donor_ sampled donor lineages S3a. The number of invisible coalescences, *n*_*IC*_, is the number of remaining recipient lineages that have not coalesced with sampled donor lineages S3b.

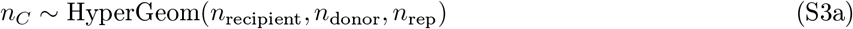

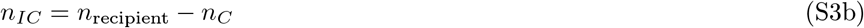

Upon a coalescence event, 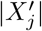 is decreased by one, a node is randomly picked in 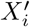, its height is updated to *t*, and it is linked to another node that is removed from 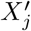. For an invisible coalescence event, 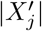 is decreased by one and both 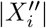 and 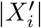 are increased by one. In this case, a new node from *X*″ is created with height *t* and linked to a node randomly picked in 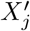. In both cases, |*X*_*i*_| is decreased by one (which is already known from the trajectory).

#### Death event

By default, a death reaction does not require any tree modification. However, if we want to simulate a full tree instead of a sampled one, the sampling events will correspond to the death events, which will therefore lead to the addition of a node. In this case, the number of sampled death events is *n*_*SD*_ = *n*_rep_.

#### Migration event

Only migration events involving deme individuals can lead to a modification in the tree. These migration reactions can be written as *X*_*i*_ → *X*_*j*_, with *j* ≠ *i*. We assume that the number of migrations that lead to a tree modification is given by the following hypergeometric distribution:

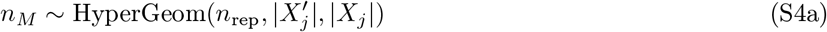

A migration increases 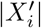 and decrease 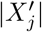 by one. A new node is created in 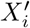 with height *t* and linked to a node randomly picked in 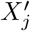, which is then removed from 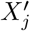. Furthermore, *X*_*i*_ is incremented by one and *X*_*j*_ is decremented by one.

### S2 Application to an epidemiological *SI*_*a*_*I*_*c*_*R* model

We illustrate the functioning of TiPS using an *SI*_*a*_*I*_*c*_*R* epidemiological compartmental model, where individuals can be susceptible (with density S), infected in acute phase (*I*_*a*_), infected in chronic phase (*I*_*c*_), and removed (R). The corresponding ODE system is

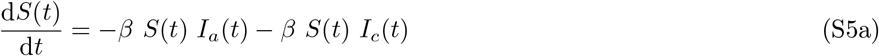

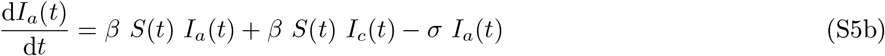

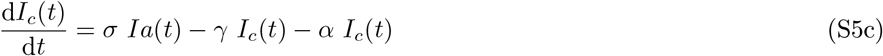

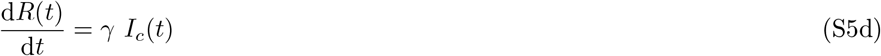

where *β* is the infectious contact rate, *γ* the recovery rate, *α* the virulence, and 1*/σ* the expected duration of the acute phase.

The model can be described as an individual-based model using a system of reactions:

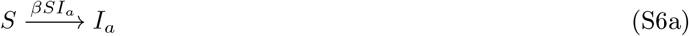

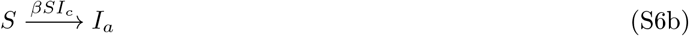

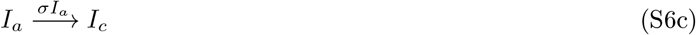

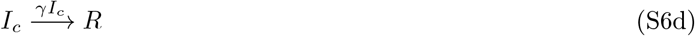

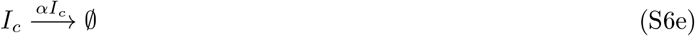

where the rate of occurrence of each reaction is indicated above the transition arrow.

Reactions S6a and S6b are birth reactions, S6c and S6d are migration reactions, and finally S6e is a death reaction event.

TiPS will first build and generate a function to simulate trajectories using this system of reactions. Population dynamics are simulated by providing parameter values using one of the algorithm described in the main text. In the following, we focus on the model captured by system of equations S6.

In this approach, under this epidemiological model, we trace back the epidemiological history of the sampled virus. Hence, the deme compartments, i.e. the ones sampled, where the virus is present and then contributing to the phylogeny, are *I*_*a*_ and *I*_*c*_.

Once the trajectory is simulated, to simulate a sampled phylogeny, TiPS requires the sampling dates and the proportion of the sampling dates to be associated with each type of deme. Let us assume, for example, that 15% of the sampling dates are associated with the *I*_*a*_ deme and 85% with the *I*_*c*_ deme compartment. TiPS randomly assigns each sampling date to a deme compartment based on these ratios and adds the dates to the list of events in the simulated trajectory (see Supplementary Figure S5). The R code to build the simulator, simulate a trajectory and a phylogeny are shown in Supplementary Figure S1.

Deme compartments (*I*_*a*_ and *I*_*c*_) can be composed of individuals represented by nodes in the simulated tree (belonging to the sub-compartments 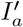 and 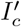). These individuals represented by nodes can also be still unsampled (belonging to 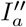 and 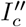).

TiPS starts the simulation of the phylogeny from the most recent sampling event and follows the trajectory backwards in time.

**Figure S1:**
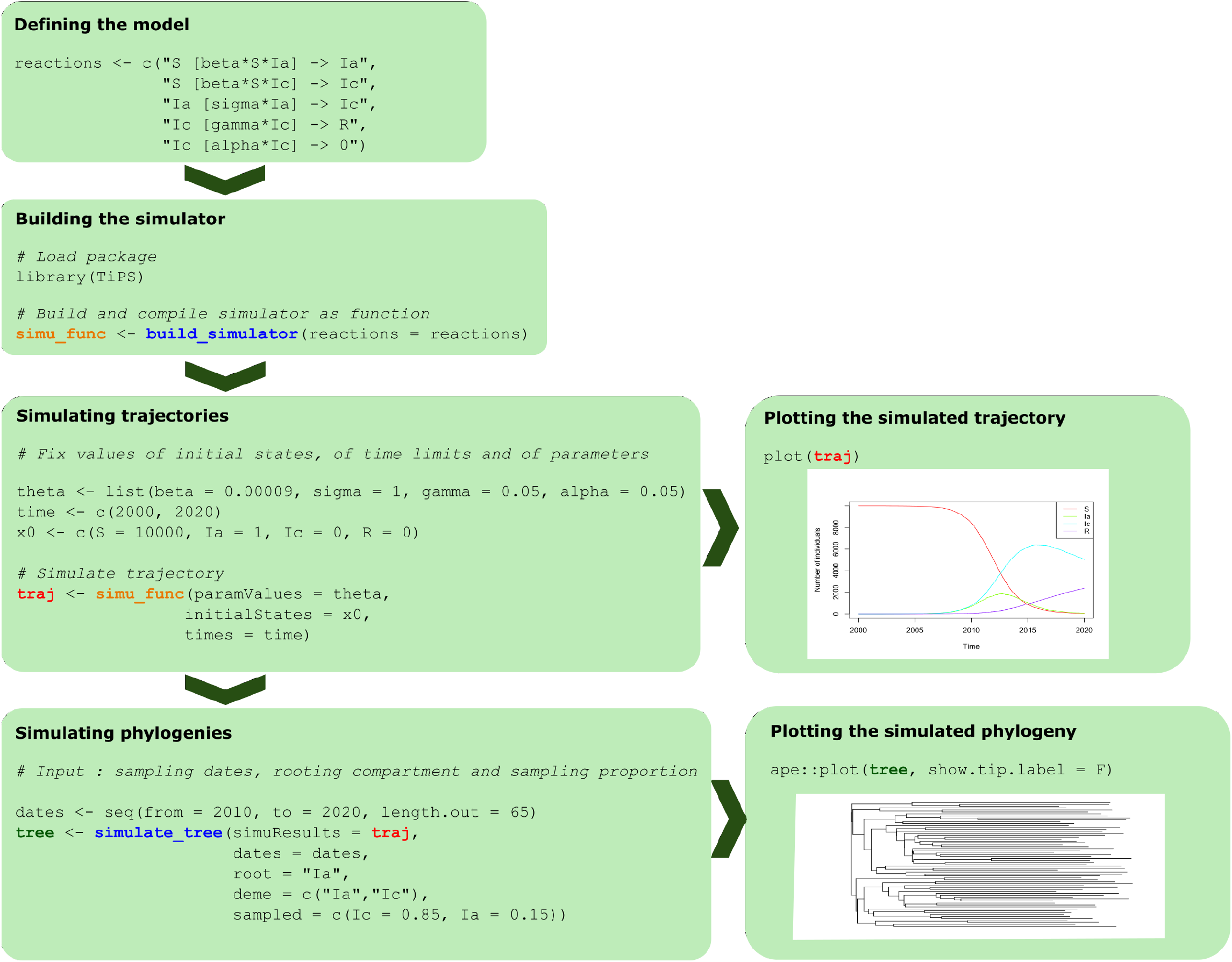
Simulating a trajectory and a phylogeny using TiPS. The equations and outputs correspond to the *SI*_*a*_*I*_*c*_*R* model. The functions of the R package are in blue. The simulator of trajectories built as a function is in orange. The variable *traj* in red is the output trajectory. The phylogeny is plotted using using the plot method of the *phylo* class. [17].

If the backward step in the trajectory leads to one or multiple migration events, i.e., in this model, from the acute phase compartment (*I*_*a*_) to the chronic infectious phase compartment (*I*_*c*_) as in reaction S6c, the number of tree modifications is given by the following hypergeometric distribution:

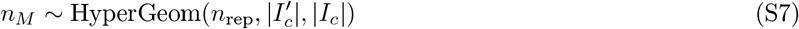

An illustration of the tree update is shown in Supplementary Figure S2.

**Figure S2:**
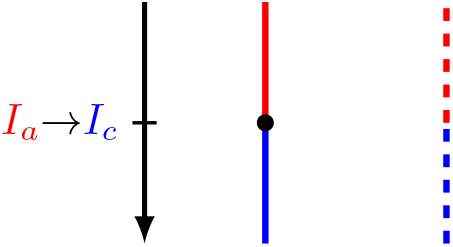
Migration event in the *SI*_*a*_*I*_*c*_*R* model. The evolution of a pathogen lineage in an individual during the acute phase of its infection is represented in red. The individual then enters the chronic phase of his infection and the pathogen lineage continues to evolve (represented in blue branches). The solid branch represents the evolution of the sampled pathogen lineage and the dashed branch represents the evolution of an unsampled pathogen lineage. The unsampled lineage is eventually removed and not represented in the final simulated phylogeny.

If the backward step in the trajectory leads to a birth reaction, two tree modifications are possible: a coalescence and an invisible coalescence. Here, we consider the donor as the host transmitting the pathogen, and the recipient the one that has been infected. In this model, there are two different demes and two different birth reactions (see reactions S6a and S6b). Given the birth reaction S6a, the number of coalescences *n*_*C*_ and invisible coalescences *n*_*IC*_ leading to tree modifications are governed by the following relationships:

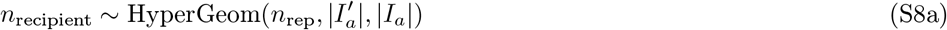

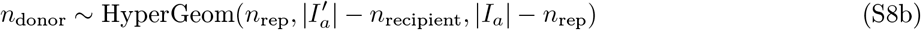

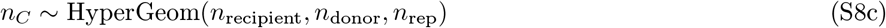

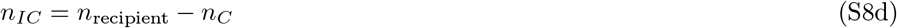

Given birth reaction S6b, the number of coalescences and invisible coalescences leading to tree modifications are governed by the following relationships:

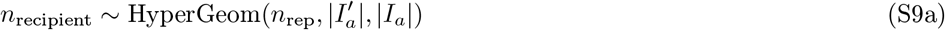

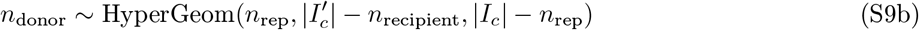

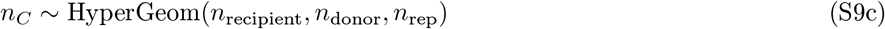

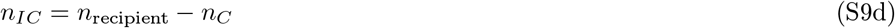

Supplementary Figure S3 illustrates possible tree updates.

This model features two types of sampling reactions: the sampling of an *I*_*a*_ individual and the sampling of an *I*_*c*_ individual. If the backward step in the trajectory leads to one or multiple sampling reactions of *I*_*a*_ individuals, the number of re-samplings 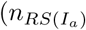 and 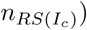 and classical samplings 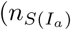 and 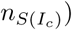 are governed by the relationships S10a and S10b, respectively. An illustration of the possible modifications in the tree are shown in Supplementary Figure S4. Note that the user can allow for re-sampling or not.

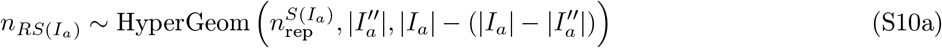

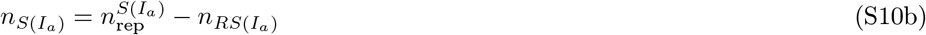

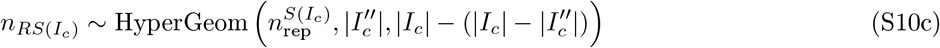

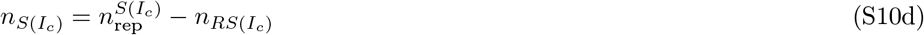

**Figure S3:**
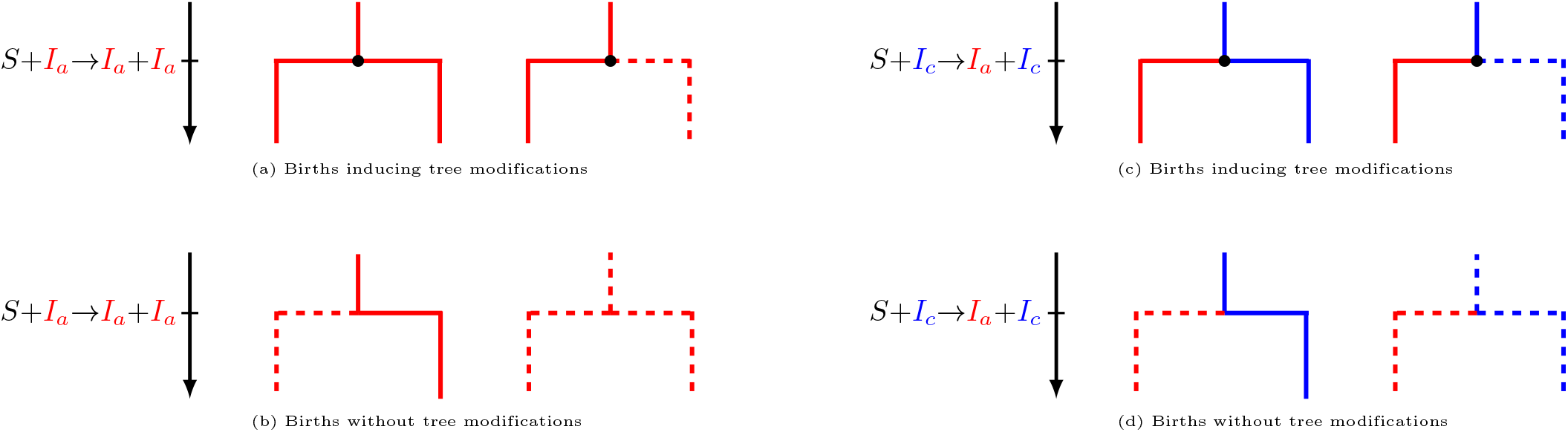
Births events in the *SI*_*a*_*I*_*c*_*R* model. Each transmission event from a living infectious individual to a susceptible individual can be represented by a branching. We show the donnor lineage on the right side of the branching and the recipient pathogen lineage (i.e. the newly-infected individual) on the left side. Dots correspond to nodes in the resulting phylogeny. Color and branch line codes are identical to Figure S2.

**Figure S4:**
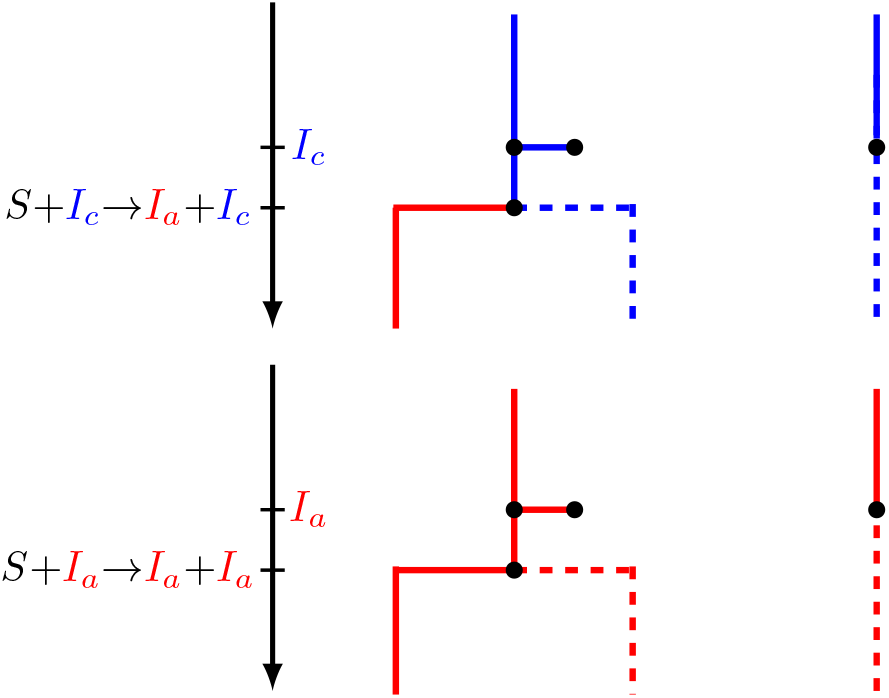
Samplings in the *SI*_*a*_*I*_*c*_*R* model. Colors, branches, and dots code are identical to Figure S3. Each sampling event leads to the addition of a node. Re-sampling events (left side of the figure) occur when the pathogen lineage has already been sampled but the individual currently carrying it has never been sampled (individuals can only be sampled once but can transmit after sampling). Otherwise, we have a classical sampling event (right side of the figure), i.e. the pathogen has not been sampled yet.

**Figure S5:**
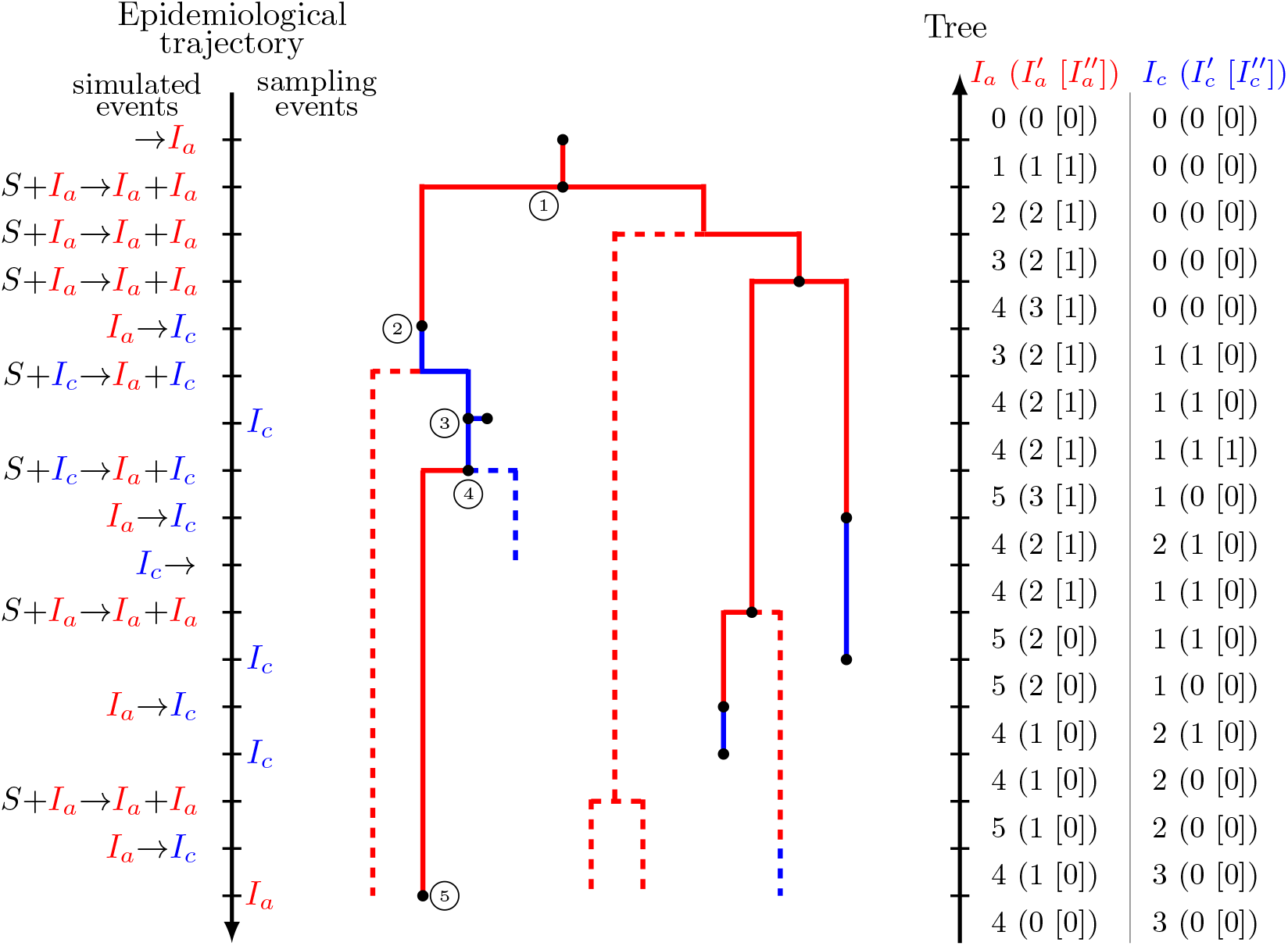
Tree simulation. We represent the epidemiological history of the individuals carrying sampled viruses in solid lines and the rest in dashed lines. Colors are identical to Figure S3. In this representation, at each birth event (branching), the donor is deviated to the right side and the recipient to the left side. All possible tree modifications are represented in this figure: (1) Coalescence; (2) Migration; (3) Re-sampling; (4) Invisible Coalescence; (5) Sampling.

### S3 Application to a logistic growth model

In this section we use the African Savannah elephant (*loxodonta africana*) population in the Kruger National Park as an example that has demonstrated an important increase in the last two decades. A study has shown that a high number of the elephants can damage the Park’s ecosystem [29]. To prevent that, until the early 90’s, a solution was the culling of elephants. Due to ethical issues, another solution has been proposed where female elephants would be on contraceptives [30].

Here we use a logistic growth model to study the population dynamics. The ODE system is:

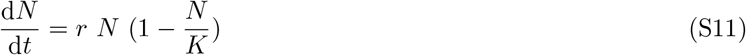

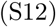

where *N* is the elephant population density, *K* is the carrying capacity of the environment, and *r* is the intrinsic growth rate of the population where *r* = *b* − *d* with *b* the birth rate and *d* the death rate.

The model can be described as an individual-based model using a system of reactions:

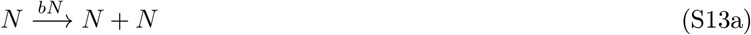

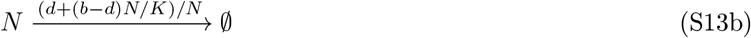

where the rate of occurrence of each reaction is indicated above the transition arrow. Reaction S13a is a birth reaction and S13b a death reaction.

TiPS allows to simulate a phylogeny without sampling dates, where events interrupting the biological process, such as deaths events or here sterilisation events, simulated in the trajectory are represented as leaves in the phylogeny. To illustrate this module of the tool, we add a sterilisation rate to the model to simulate the events in the trajectory. The system of reactions becomes:

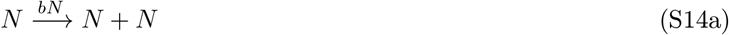

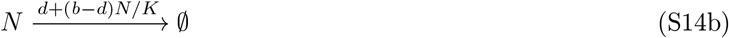

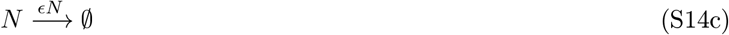

where reaction S14c is the sterilisation reaction with a rate of its occurrence indicated above the transition error. Using this system of reactions, TiPS will first build and generate a function to simulate trajectories. Population dynamics are simulated by providing parameter values using one of the algorithm described in the main text. The R code to build the simulator, simulate a trajectory and a phylogeny are shown in Supplementary Figure S6.

We assume that a sampling event is an event that interrupts a biological process. In this example, where no sampling dates are required, the death events and the elephant sterilisation events will be considered as the sampling events and will be represented as leaves in the simulated phylogeny.

We introduce here the compartments *N, N*′ and *N*′′ where *N*′′ *⊆ N*′ *⊆ N*. All the elephants are in compartment *N*, the sampled elephants are sub-compartment *N* and the elephants that in *N* that have not yet been but may be sampled are in sub-compartment *N*′′.

TiPS starts the simulation of the phylogeny from the most recent sampling event and follows the trajectory backwards-in-time.

When the backward step in the trajectory at time *t* leads to *n* death or sterilisation events, *n* new nodes are created with height *t* in *N*′ and |*N*′ | increases by one. Note that, each node is labelled with the corresponding reaction, so we can distinguish and visualise them when plotting the phylogeny.

If the backward step in the trajectory leads to a birth reaction, two tree modifications are possible: a coalescence and an invisible coalescence. A coalescence in the phylogeny corresponds to the coalescence of two sampled lineages each representing an elephant, into one sampled individual (the donor). An ‘invisible’ coalescence is the coalescence of the sampled lineage of the recipient individual (*N*′) and an unsampled lineage of the donor individual (*N*′′) into one unsampled lineage (the donor, *N*′′).

Given the birth reaction S14a, the number of coalescences *n*_*C*_ and invisible coalescences *n*_*IC*_ leading to tree modifications are governed by the following relationships:

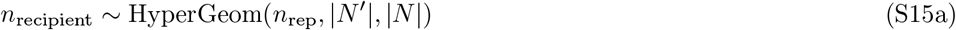

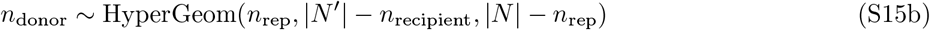

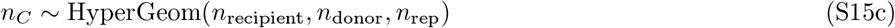

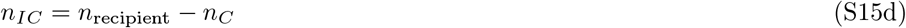

where *n*_*rep*_ is the number of birth events in the trajectory at time t of the trajectory, *n*_*recipient*_ is the number of recipient lineages and *n*_*donor*_ the number of donor lineages. Upon a coalescence, |*N*′ | is decreased by one, a node is randomly picked from *N*′ with height *t* and is linked to another node that is removed from *N*′. For an invisible coalescence, |*N*′ | is decreased by one both and *N*′′ and *N*′ are increased by one. A new node from *N*′′ is created with height *t* ans is linked to a node randomly picked in *N*′. In both cases, |*N* | is decreased by one as recorded already in the simulated trajectory.

**Figure S6:**
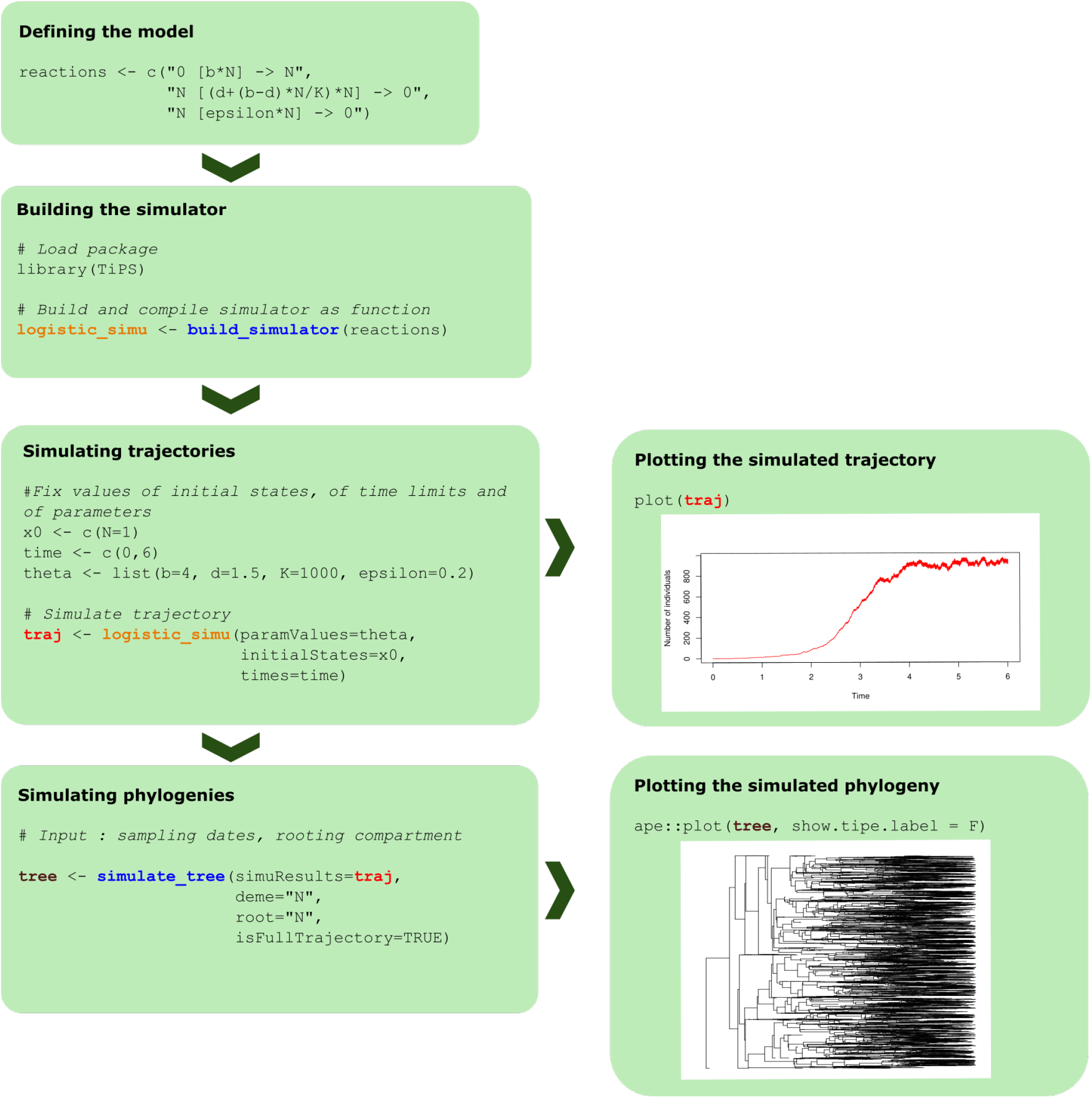
Simulating a trajectory and a phylogeny using TiPS given a logistic growth model. The equations and outputs correspond to the logistic growth model. The functions of the R package are in blue. The simulator of trajectories built as a function is in orange. The variable *traj* in red is the output trajectory. The phylogeny is plotted using a function from the ape R package. [17].

### S4 Benchmarking

#### S4.1 Benchmarking: trajectories

To evaluate our simulator we performed a benchmarking analysis on both modules of the tool, i.e. the trajectory and the phylogeny simulators, using two existing `R` packages. `adpativetau` performs simulations of trajectories of continuous-time Markov processes by using Gillespie’s stochastic simulation algorithms. `phydynR` is a package for phylodynamic inference using population genetic models and performs simulations of trajectories as well using the Euler-Maruyama integration method and provides methods for simulating trees conditional on a demographic process. The different algorithms of each package are presented in Table S1.

**Supplementary Table S1:**
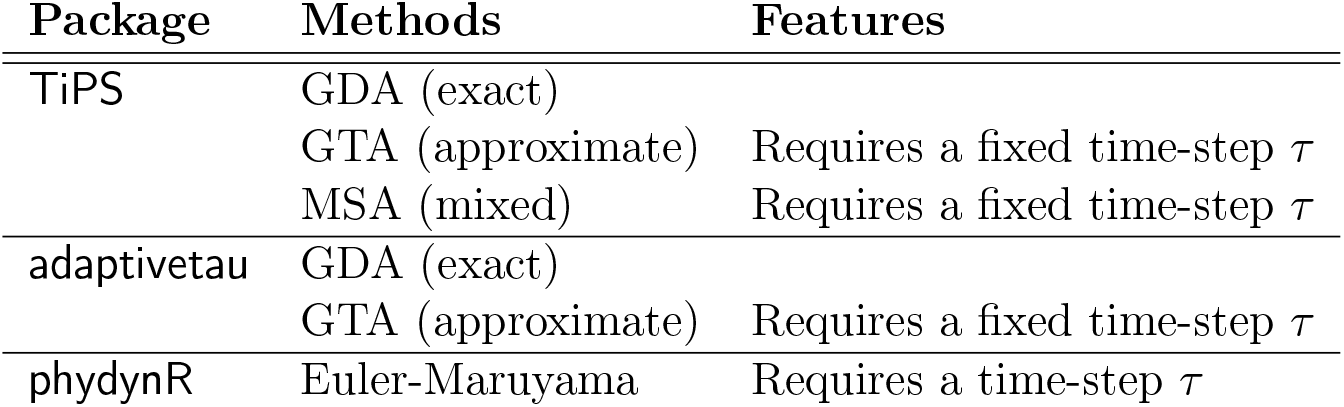
Main algorithms of the R packages compared. GDA stands for Gillespie’s Direct Algorithm, GTA for Gillespie’s Tau-Leap Algorithm, and MSA for Mixed simulation algorithm.

#### S4.2 Benchmarking: phylogenies

To evaluate the accuracy of phylogeny simulations, we generated 10 different sub-trees of 1, 000 leaves, using detailed epidemiological model with two host types, as described in [26]. We then used `TiPS` and `phydynR` to simulate 1, 000 phylogenies with each package under the same epidemiological model with the same parameter values, imposing the 1, 000 dates of each target sub-tree under a backwards-in-time approach. To compare the simulated phylogenies with the target one, we computed summary statistics for each of them using the methods described in [9]. Supplementary Figure S7 shows the distributions of summary statistics computed from the phylogenies simulated using `TiPS` (in red) and using `phydynR` (in orange) for each analysis using a different target tree (in black).

**Figure S7:**
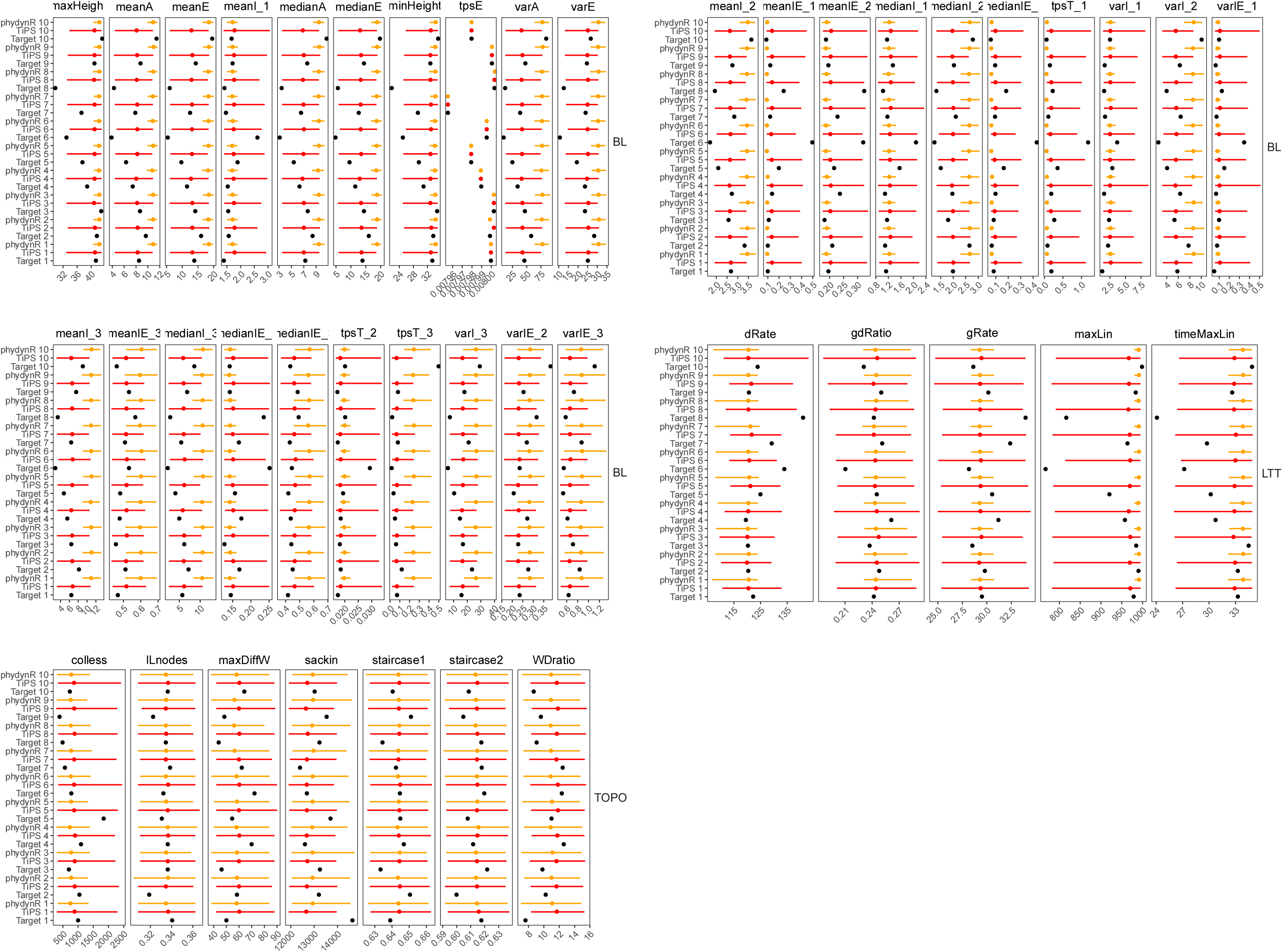
Distributions of summary statistics. The summary statistics are grouped into three families : branch lengths (denoted BL), tree topology (TOPO) and Lineage-Throuht-Time plot (LTT). The dots represent the median and the horizontal lines represent the 95% HPD. Red distributions correspond to the summary statistics computed from the phylogenies simulated using `TiPS`, orange distributions correspond to the summary statistics computed from the phylogenies simulated using `phydynR`, for each target tree analysis. Black dots represent the values of summary statistics computed from each target tree.

## References

[1] Otto SP, Day T. A biologist’s guide to mathematical modeling in ecology and evolution. Princeton University Press; 2007.

[2] Keeling MJ, Rohani P. Modeling infectious diseases in humans and animals. Princeton University Press; 2008.

[3] Lenormand T, Roze D, Rousset F. Stochasticity in evolution. Trends in Ecology and Evolution. 2009;24(3):157– 165.

[4] Grenfell BT, Pybus OG, Gog JR, Wood JLN, Janet M, Mumford JA, et al. Unifying the Epidemiological and Evolutionary Dynamics of Pathogens. Science. 2004;303(5656):327–332.

[5] Volz EM, Koelle K, Bedford T. Viral phylodynamics. PLoS Comput Biol. 2013;9(3):e1002947.

[6] Frost SD, Pybus OG, Gog JR, Viboud C, Bonhoeffer S, Bedford T. Eight challenges in phylodynamic inference. Epidemics. 2015;10:88–92.

[7] Ratmann O, Donker G, Meijer A, Fraser C, Koelle K. Phylodynamic inference and model assessment with approximate bayesian computation: influenza as a case study. PLoS Comput Biol. 2012;8(12):e1002835.

[8] Gascuel F, Ferrière R, Aguilée R, Lambert A. How Ecology and Landscape Dynamics Shape Phylogenetic Trees. Systematic Biology. 2015;64(4):590–607.

[9] Saulnier E, Gascuel O, Alizon S. Inferring epidemiological parameters from phylogenies using regression-ABC: A comparative study. PLOS Computational Biology. 2017 Mar;13(3):e1005416.

[10] Gillespie DT. A general method for numerically simulating the stochastic time evolution of coupled chemical reactions. Journal of Computational Physics. 1976;22(4):403–434.

[11] Kurtz TG. Solutions of ordinary differential equations as limits of pure jump markov processes. Journal of Applied Probability. 1970;7(1):49–58.

[12] Pineda-Krch M. GillespieSSA: implementing the stochastic simulation algorithm in R. Journal of Statistical Software. 2008;25(12):1–18.

[13] Johnson P. adaptivetau: efficient stochastic simulations in R. R Package Version. 2014;.

[14] Bjørnstad ON. Epidemics. Models and Data using R. Springer; 2018.

[15] Pennell M, Eastman J, Slater G, Brown J, Uyeda J, Fitzjohn R, et al. geiger v2.0: an expanded suite of methods for fitting macroevolutionary models to phylogenetic trees. Bioinformatics. 2014;30:2216–2218.

[16] Revell LJ. phytools: An R package for phylogenetic comparative biology (and other things). Methods in Ecology and Evolution. 2012;3:217–223.

[17] Paradis E, Schliep K. ape 5.0: an environment for modern phylogenetics and evolutionary analyses in R. Bioinformatics. 2019;35:526–528.

[18] Volz EM. Complex population dynamics and the coalescent under neutrality. Genetics. 2012;190(1):187–201.

[19] Vaughan TG, Drummond AJ. A stochastic simulator of birth-death master equations with application to phylodynamics. Molecular Biology and Evolution. 2013;30(6):1480–1493.

[20] Bouckaert R, Heled J, Kühnert D, Vaughan T, Wu CH, Xie D, et al. BEAST 2: A Software Platform for Bayesian Evolutionary Analysis. PLoS Computational Biology. 2014;10(4):1–6.

[21] Eddelbuettel D, Francois R. Rcpp: Seamless R and C++ Integration. J Stat Software. 2011;40(1):1–18.

[22] Gillespie DT. Approximate accelerated stochastic simulation of chemically reacting systems. Journal of Chemical Physics. 2001;115(4):1716–1733.

[23] Cao Y, Gillespie DT, Petzold LR. The slow-scale stochastic simulation algorithm. The Journal of Chemical Physics. 2005;122(1):014116. Available from: https://doi.org/10.1063/1.1824902.

[24] Cao Y, Gillespie DT, Petzold LR. Adaptive explicit-implicit tau-leaping method with automatic tau selection. Journal of Chemical Physics. 2007;126(22):1–10.

[25] Kingman JFC. The coalescent. Stochastic processes and their applications. 1982;13(3):235–248.

[26] Danesh G, Virlogeux V, Ramière C, Charre C, Cotte L, Alizon S. Quantifying transmission dynamics of acute hepatitis C virus infections in a heterogeneous population using sequence data. PLoS Pathogens. 2021;17(9):e1009916.

[27] Hartfield M, Alizon S. Epidemiological feedbacks affect evolutionary emergence of pathogens. Am Nat. 2014;183(4):E105–E117.

[28] Alizon S. Superspreading genomes. Science. 2021;371(6529):574–575.

[29] Cumming DH, Fenton MB, Rautenbach IL, Taylor RD, Cumming GS, Cumming MS, et al. Elephants, woodlands and biodiversity in southern Africa. South African Journal of Science. 1997;93(5):231–236.

[30] Whyte I, van Aarde R, Pimm SL. Managing the elephants of Kruger National Park. Animal Conservation. 1998;1(2):77–83.

